# High aspect ratio graphene oxide: a highly efficient plasmid DNA deliverer for plant seed

**DOI:** 10.1101/2024.10.06.616812

**Authors:** Cheng Jiang, Yue Pan, Xinyu Li, Zhongzhu Yang

## Abstract

Plant genetic engineering plays a central role in crop improvement and the biosynthesis of natural products. However, the plant cell wall serves as a natural barrier that restricts the efficient delivery of exogenous biomolecules, particularly the direct transfer of plasmid DNA into plant seeds. Here, we introduce a novel form of graphene oxide characterized by a high aspect ratio, synthesized through low-voltage, low-current, and prolonged electrochemical oxidation in a 0.5 mol/L NaOH aqueous solution. The high aspect ratio graphene oxide (HARGO) can effectively deliver plasmid DNA to *Poa crymophila* Keng seeds, achieving a 90% success rate. Furthermore, it is effective for both wheat and rice seeds. The underlying mechanism of this efficient delivery is that HARGO can physically absorb plasmid DNA and transport the adsorbed plasmid DNA into plant cells and deeper tissues through passive transport, eliminating the need for chemical modification of polyethylene glycol (PEG) and polyethyleneimine (PEI). These findings offer a dependable non-integrating plasmid DNA delivery tool for plant genetic engineering, which will significantly impact the advancement of plant biotechnology.

---

Plant biotechnology plays a crucial role in addressing global food security and energy demands, as well as having significant applications in the field of biopharmaceuticals^1,2,3^. In agriculture, genetic improvement techniques can develop varieties that are high-yielding, high-quality, resistant to herbicides, diseases, and pests, as well as tolerant to abiotic stresses^4^. In the medical field, genetically engineered plants can produce valuable microbial drugs and recombinant proteins, while bioengineered plants can also generate cleaner and more efficient biofuels^5,6,7^.

Although plant biotechnology has made numerous advancements over the past few decades, the genetic transformation of many plant species remains highly challenging^8^. One of the primary challenges is to penetrate the tough and multi-layered plant cell walls to effectively deliver biomolecules into the cells and even deeper into the tissues^8,9^. Currently, only a limited number of delivery tools are employed to achieve this goal, each with its own constraints. *Agrobacterium*-mediated delivery is the most commonly used method for plant gene delivery, but its effectiveness is limited to specific plant species and tissue types, and it is not suitable for DNA-free and foreign gene-free editing processes^10^. The biolistic particle delivery system, also known as the gene gun, is another commonly used tool for plant transformation. It can deliver bio-molecules to a broader range of plants but has several drawbacks, including expression only at the point of impact, potential damage to plant tissues, limitations on sample size, the need for precise targeting, and the requirement of large amounts of DNA for efficient delivery^11^. For the transient expression of foreign proteins in plants, technologies based on the tobacco mosaic virus, such as Geneware, and other plant viral vectors like the potato X virus and cowpea mosaic virus, facilitate the industrial-scale production of relevant proteins. However, these viral vectors have limited compatibility with specific plant species and the size of the expression vector, which restricts the choice of host plants and hinders the simultaneous expression of large proteins or multiple proteins. Furthermore, the use of viral vectors for transient expression in gene editing systems is typically subject to stringent regulatory scrutiny due to their pathogenic origins and the potential for some viruses to integrate their genetic material into the plant genome^12,13,14^. Therefore, to address these challenges, researchers are actively exploring new technologies and methods to enhance the efficiency and applicability of plant genetic transformation.

The application of nanomaterials for gene delivery to animal cells has been extensively researched; however, their potential in plant systems remains underexplored. While research indicates that plant cells can absorb nanomaterials, many of these investigations focus primarily on the delivery of non-functional payloads or depend on mechanical methods—such as gene guns or ultrasound—to promote the penetration of nanoparticles through the plant cell wall.^15^. Recently, several nanomaterials, such as mesoporous silica nanoparticles (MSNs), DNA nanostructures, silicon carbide whiskers (SCWs), layered double hydroxide (LDH) clay nanosheets, and carbon nanotubes (CNTs), have shown the potential to penetrate the plant cell wall and deliver functional biological payloads without the need for strong mechanical assistance. In the study of MSNs, MSNs have been shown to efficiently deliver plasmid DNA to *Arabidopsis* roots and deliver siRNA to leaves of mature plants through passive transport^16,17^. SCWs have enabled the delivery of genes to undifferentiated plant tissues and explants in suspension by co-culturing and vertexing with plant cells and DNA, promoting the stable transformation and screening of transgenic plants. It is speculated that SCWs may allow DNA entry into cells by penetrating or tearing the cell wall, but this mechanism is not suitable for subcellular/tissue targeting or testing in whole plants, and may affect transformation efficiency and cell health^8,18^. LDH clay nanosheets have effectively delivered RNAi molecules to tobacco, achieving gene silencing and providing a new direction for the development of plant bio-nanotechnology. Although LDH clay nanosheets have not yet been used for plasmid DNA delivery, their potential is promising^19^. CNTs, emerging as carriers for plant plasmid DNA delivery, have demonstrated the ability to effectively adsorb and deliver plasmid DNA to plant cells, resulting in the expression of fluorescent proteins. However, the size of chemically modified CNTs increases significantly after binding to plasmid DNA, restricting their entry to either plant protoplasts or the surface cells of plant tissue^20,21,22^. In summary, the prospects for the application of nanomaterials in plant gene delivery are extensive; however, challenges remain, including tissue penetration, cell health, and subcellular targeting, necessitating further research and optimization.

Graphene oxide (GO) is an oxidized form of graphene characterized by a substantial presence of oxygen-containing functional groups, mainly including carboxyl, carbonyl, and epoxy groups. GO exhibits properties such as a large surface area, strong hydrophilicity, and low biological toxicity^23^. GO is typically synthesized from graphite through strong acid oxidation employing methods such as Brodie’s method, Staudenmaier’s method, or Hummers’ method^24^. Existing research indicates that GO prepared by traditional methods can not only enter plant cells via passive transport but also bind with double-stranded DNA under specific ionic conditions^25^. Currently, using GO as a carrier for delivering plasmid DNA into plant cells necessitates modification with PEG and PEI. Similar to CNTs, chemically modified GO increases significantly in size after adsorbing plasmid DNA, thereby restricting delivery to plant protoplasts or the surface cells of plant tissues^26^. To date, there are no reports of utilizing chemically unmodified GO for delivering plasmid DNA to plant seeds.

To address the limitations of existing techniques, this study developed a novel type of GO with a high aspect ratio (high aspect ratio graphene oxide, HARGO) through low-voltage, low-current, and prolonged electrochemical oxidation in dilute NaOH aqueous solution. Unmodified HARGO can physically adsorb plasmid DNA in aqueous solution and demonstrates potential for penetrating the rigid cell walls of plants, reaching the embryonic portions of seeds, and facilitating the formation of positively transformed seedlings. This innovative approach offers new opportunities to enhance the efficiency and applicability of plant genetic transformation.

## Preparation and characterization of HARGO

Electrolysis was conducted using a graphite rod (5 mm in diameter) as the anode and a platinum (Pt) electrode as the cathode in a 0.5 M NaOH aqueous solution for 96 hours at a voltage of 3.5 V and a current of 0.03 A. The GO mixed aqueous solution was successfully synthesized. The resulting mixed aqueous solution is brownish, uniformly dispersed, and exhibits no noticeable precipitation (Fig. 1a). Following dialysis and purification, GO powder was obtained through drying. The GO powder was dissolved in water, and following ultrasonic dispersion, GO aqueous solutions with concentrations of 10, 5, and 1 mg/mL were prepared (Fig. 1a). To observe the microstructural characteristics of GO prepared by this method, atomic force microscopy (AFM) analysis was performed. The AFM results indicate that the lamellar thickness of GO prepared by this method is 0.414 nm. The morphology is elongated, with a length-to-width ratio of approximately 8:1, and it exhibits longitudinal cracks along the long axis (Fig. 1b). Given the unique high aspect ratio morphology and structural characteristics of this GO, which differ from those of ordinary GO, it is designated as high aspect ratio GO (HARGO).

**Fig. 1.**
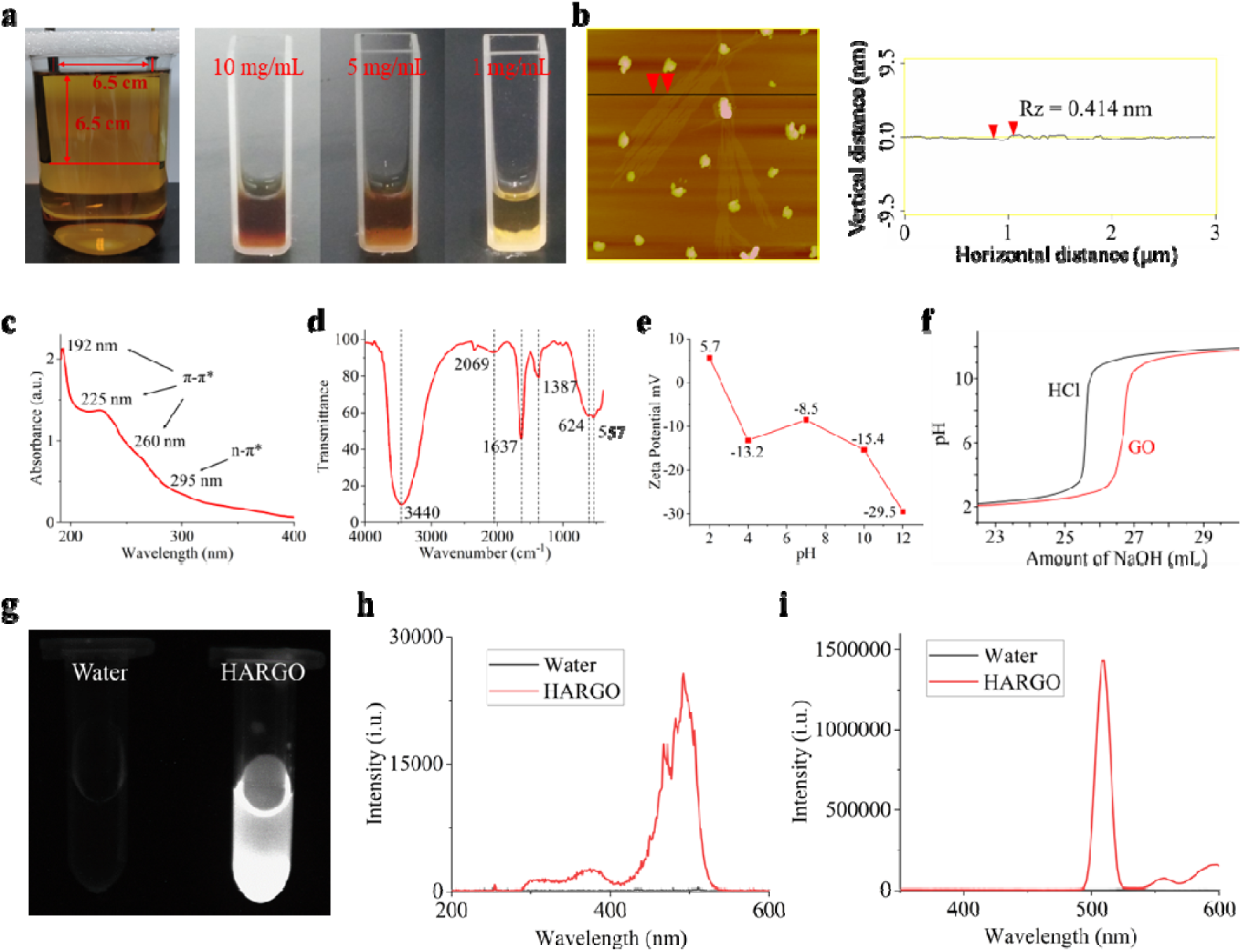
Preparation and characterization of high aspect ratio graphene oxide (HARGO). **a**, The left image displays a digital photograph taken after 96 hours of electrolysis, in which the anode is a graphite rod and the cathode is a Pt electrode, positioned 6.5 cm apart and submerged 6.5 cm below the liquid surface. The right image illustrates HARGO aqueous solutions at concentrations of 10, 5, and 1 mg/mL, respectively. **b**, The left image presents an atomic force microscope (AFM) image of HARGO, with the point-like substance identified as NaOH impurity. The right image depicts the height profile. **c**, UV-vis Spectrum of HARGO at a concentration of 0.5 mg/mL. **d**, Fourier transform infrared spectrum of HARGO. **e**, Zeta Potential Analysis of HARGO at a concentration of 0.1 mg/mL. **f**, Titration curve of HARGO. **g**, Digital photograph of fluorescence emitted by 1 mg/mL HARGO aqueous solution under excitation light with a wavelength of 350 nm. **H**, Excitation spectrum of HARGO. **i**, Emission spectrum of HARGO.

Subsequently, HARGO was further characterized. UV-Vis analysis revealed a prominent absorption peak at 192 nm, along with three shoulder peaks at 225 nm, 260 nm, and 295 nm in the aqueous solution of HARGO. The absorption peaks at 192 nm, 225 nm, and 260 nm are specifically attributed to the characteristic π-π* transitions of HARGO, whereas the peak at 295 nm corresponds to n-π* transitions (Fig. 1c). Fourier transform infrared spectrum (FT-IR) analysis identified absorption peaks for HARGO at 3440 cm^-1^, 2069 cm^-1^, 1637 cm^-1^, 1387 cm^-1^, 624 cm^-1^, and 557 cm^-1^. The absorption peaks at 1637 cm_¹ and 1387 cm_¹ indicate the presence of carboxyl groups (COO_), while the peak at 3440 cm_¹ signifies the existence of hydroxyl groups (-OH). The absence of a peak at 1720 cm_¹ suggests a lack of free carbonyl groups at the edges (Fig. 1d). Zeta potential test results indicate that the content of oxygen-containing groups per unit area on the surface of HARGO (pH = 7, ζ = -8.5 mV) is lower than that of GO prepared by the traditional Hummers method (pH = 7, ζ = -27.5 mV) (Fig. 1e). Titration test results indicate that the pKa of the carboxyl group in HARGO is approximately 4.82, the concentration of the carboxyl group is about 7.8 × 10_³ mol/g, and the surface charge per unit mass is approximately -751 C/g, which is significantly higher than that of GO prepared by the traditional Hummers method (Fig. 1f)^24^.

Surprisingly, when ultraviolet light irradiates the HARGO aqueous solution, it is observed that HARGO emits fluorescence (Fig. 1g). Subsequently, the excitation and emission spectra of the HARGO aqueous solution were analyzed. The excitation spectrum indicates that the maximum excitation wavelength of HARGO is 492 nm, with an excitation wavelength range of approximately 290-530 nm (Fig. 1h). The emission spectrum reveals that the maximum emission wavelength of HARGO is 510 nm (Fig. 1i).

## HARGO can significantly penetrate various tissues and cells in wheat seeds

By utilizing the fluorescent properties of HARGO, we evaluated its ability to penetrate plant seeds. We used spring wheat Zhongkemai 138 (ZKM138) seeds and soaked them in a HARGO aqueous solution for 24 hours before performing frozen sectioning. To ensure that the plant cells were alive during observation, we examined all samples immediately after frozen sectioning. Inverted fluorescence microscopy results indicated that a substantial amount of HARGO was detected in the groin, endosperm, germ, and germ base of wheat seeds (Fig. 2a). Further analysis using laser confocal microscopy revealed that HARGO was abundantly distributed in the germ cambium, establishing a basis for delivering plasmid DNA to plant seeds and subsequently generating positive seedlings (Fig. 2b). To investigate whether HARGO penetrates the interior of plant cells, the roots of wheat seedlings soaked in HARGO aqueous solution were subjected to frozen sectioning and microstructural analysis. Inverted fluorescence microscopy revealed a substantial amount of HARGO in the roots, and laser confocal optical sectioning demonstrated that HARGO was not only significantly enriched on the cell wall but also present in large quantities within the cells (Fig. 2c). To more intuitively illustrate the process of HARGO penetrating plant cells, we created a schematic diagram depicting HARGO’s passive transport into plant cells based on the experimental results (Fig. 2d).

**Fig. 2.**
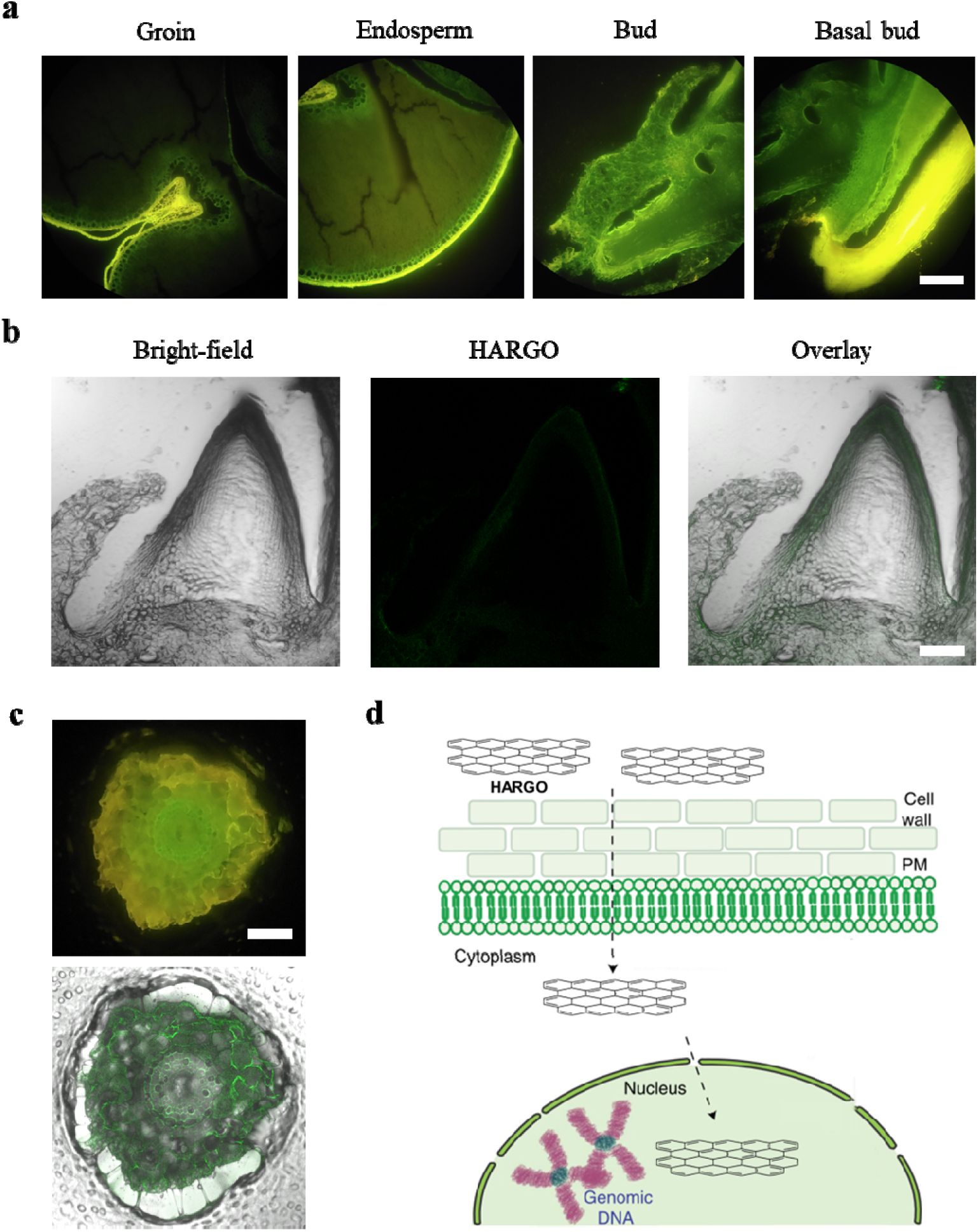
The distribution of high aspect ratio graphene oxide (HARGO) in various tissues and cells of wheat. **a**, Inverted fluorescence microscopy images of frozen sections of spring wheat Zhongkemai 138 (ZKM138) seeds, which were soaked in a 0.5 mg/mL HARGO aqueous solution for one day, depict the groin, endosperm, bud, and basal part of the embryo from left to right. The scale bar represents 25 µm. **b**, Laser confocal images of the bud are presented from left to right as bright field, HARGO, and overlay. The scale bar represents 25 µm. **c**, Inverted fluorescence microscopy image (top) and laser confocal microscopy image (bottom) of frozen cross-sections of wheat roots after 14 days post-seedling emergence. The scale bar represents 25 µm. **d**, Schematic representation of HARGO penetrating plant cells.

## Adsorption Capacity of HARGO for Plasmid DNA pEG100-PcNAC-EGFP

Given HARGO’s excellent penetrability into plant tissues, it is essential for HARGO to bind to plasmid DNA for its effective transportation. To investigate HARGO’s adsorption capacity for plasmid DNA, we first constructed the plasmid DNA pEG100-PcNAC-EGFP, which is 11,798 bp in length (Fig. S1). Subsequently, we established four experimental groups: a water-only control group, a HARGO-only group, a pEG100-PcNAC-EGFP-only group, and a mixed group containing both HARGO and pEG100-PcNAC-EGFP. UV-Vis analysis revealed that in the mixed group of HARGO and pEG100-PcNAC-EGFP, the absorbance value at 225 nm was 16.24, significantly lower than the combined absorbance values of the HARGO-only group (16.11) and the pEG100-PcNAC-EGFP-only group (0.49), resulting in an expected value of 16.60. Furthermore, compared to the HARGO-only group, the mixed group exhibited a redshift in the absorbance peak at 225 nm, suggesting that HARGO may have physically adsorbed plasmid DNA (Fig. 3a).

**Fig. 3.**
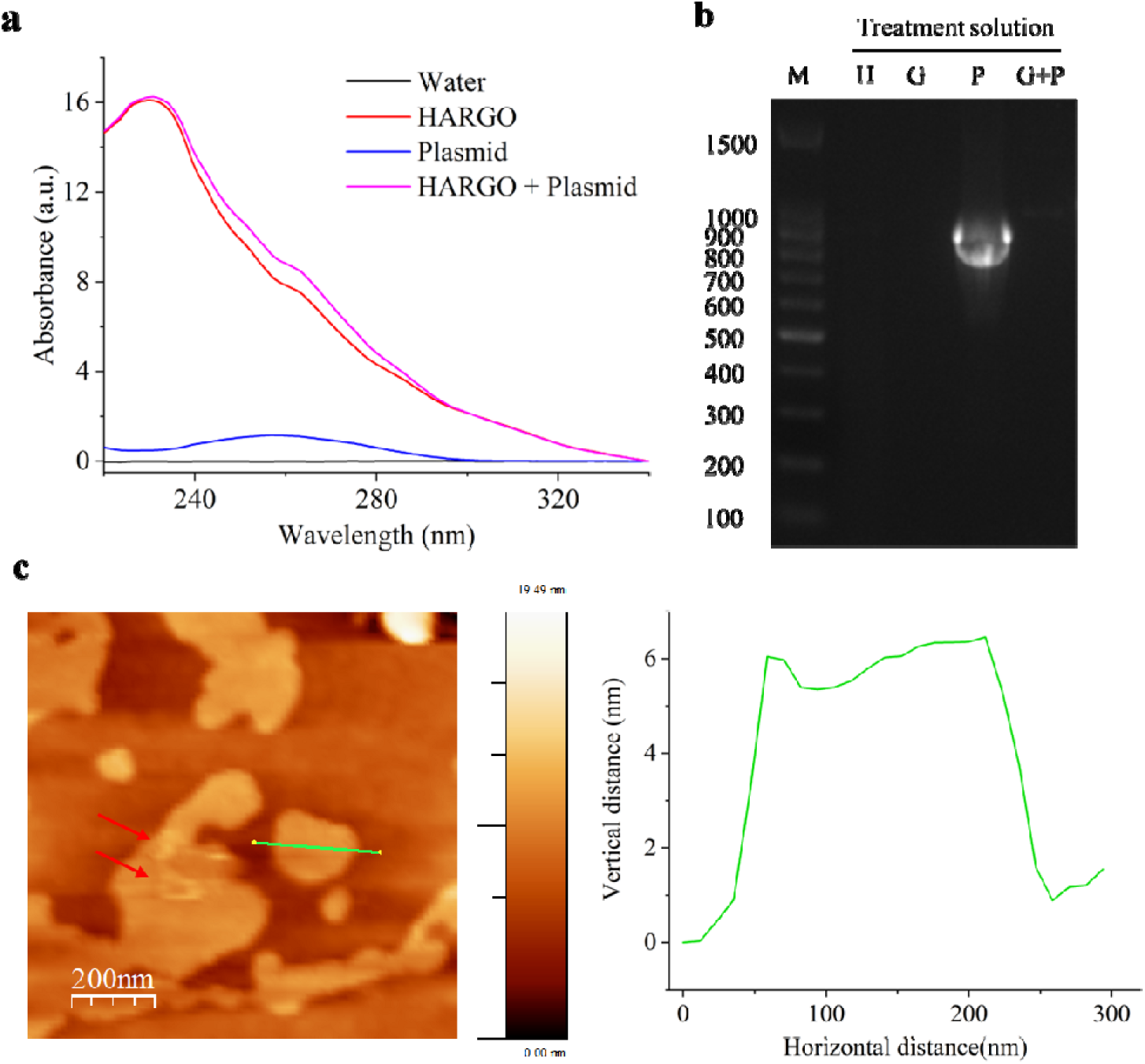
High aspect ratio graphene oxide (HARGO) adsorbs plasmid DNA pEG100-PcNAC-EGFP. **a**, UV spectra of water, HARGO, plasmid DNA pEG100-PcNAC-EGFP, and mixed aqueous solutions of HARGO and pEG100-PcNAC-EGFP. **b**, Digital photographs of agarose gel electrophoresis after PCR amplification were obtained using water (H), HARGO (G), plasmid DNA pEG100-PcNAC-EGFP (P), and their mixed aqueous solution (G+P) as templates. **c**, The left image displays an atomic force microscopy image of a mixed aqueous solution of HARGO and pEG100-PcNAC-EGFP, with arrows indicating the presence of pEG100-PcNAC-EGFP. The right image depicts the height profile.

To confirm the physical adsorption of HARGO to pEG100-PcNAC-EGFP, a PCR analysis was conducted. The results indicated that the mixture of HARGO and pEG100-PcNAC-EGFP did not amplify the target DNA band, suggesting that physical adsorption occurred between HARGO and the plasmid DNA. This adsorption impeded the attachment of DNA polymerase to the plasmid DNA adsorbed on the HARGO surface (Fig. 3b). Lastly, we employed AFM to assess the status of HARGO adsorbing pEG100-PcNAC-EGFP. The results clearly confirmed that HARGO can adsorb pEG100-PcNAC-EGFP (Fig. 3c).

## HARGO-mediated delivery of pEG100-PcNAC-EGFP plasmid DNA to *Poa crymophila* Keng, wheat, and rice seeds

The results of the research indicate that HARGO exhibits high penetrability, allowing it to penetrate various tissues and cells of wheat while also physically adsorbing plasmid DNA. Based on these findings, we hypothesize that HARGO may possess significant potential for delivering plasmid DNA into plant seeds.

The low or negligible toxicity of HARGO to plants is a critical factor for its application in plant genetic transformation. Consequently, we initially evaluated the biological toxicity of HARGO. Using spring wheat ZKM138 as the experimental material, we co-cultured ZKM138 seeds with a HARGO aqueous solution at a concentration of 0.5 mg/mL. The results indicate that HARGO is not only non-toxic to ZKM138 but also significantly enhances the growth of both seedlings and roots (Fig. S2a-d). Additionally, HARGO does not significantly affect the seedling emergence rate can significantly reduce the number of moldy seeds, suggesting that it may possess an ability to inhibit mold growth (Fig. S2c).

Building on this premise, we conducted a study to further investigate whether HARGO can effectively deliver plasmid DNA to plant seeds, using *Poa crymophila* Keng cv. Qinghai seeds as the experimental material. The *Poa crymophila* Keng cv. Qinghai seeds were sown in 2-milliliter Eppendorf tubes containing a dry matrix soil at a density of 20 seeds per tube and irrigated with water, HARGO aqueous solution, plasmid DNA pEG100-PcNAC-EGFP aqueous solution, and a HARGO + pEG100-PcNAC-EGFP mixed aqueous solution, respectively. After 14 days of seedling emergence, we observed the seedlings (Fig. 4a), and the statistical analysis of seedling height indicated that both the HARGO aqueous solution and the HARGO + pEG100-PcNAC-EGFP mixed aqueous solution significantly promoted seedling growth (Fig. 4b).

**Fig. 4.**
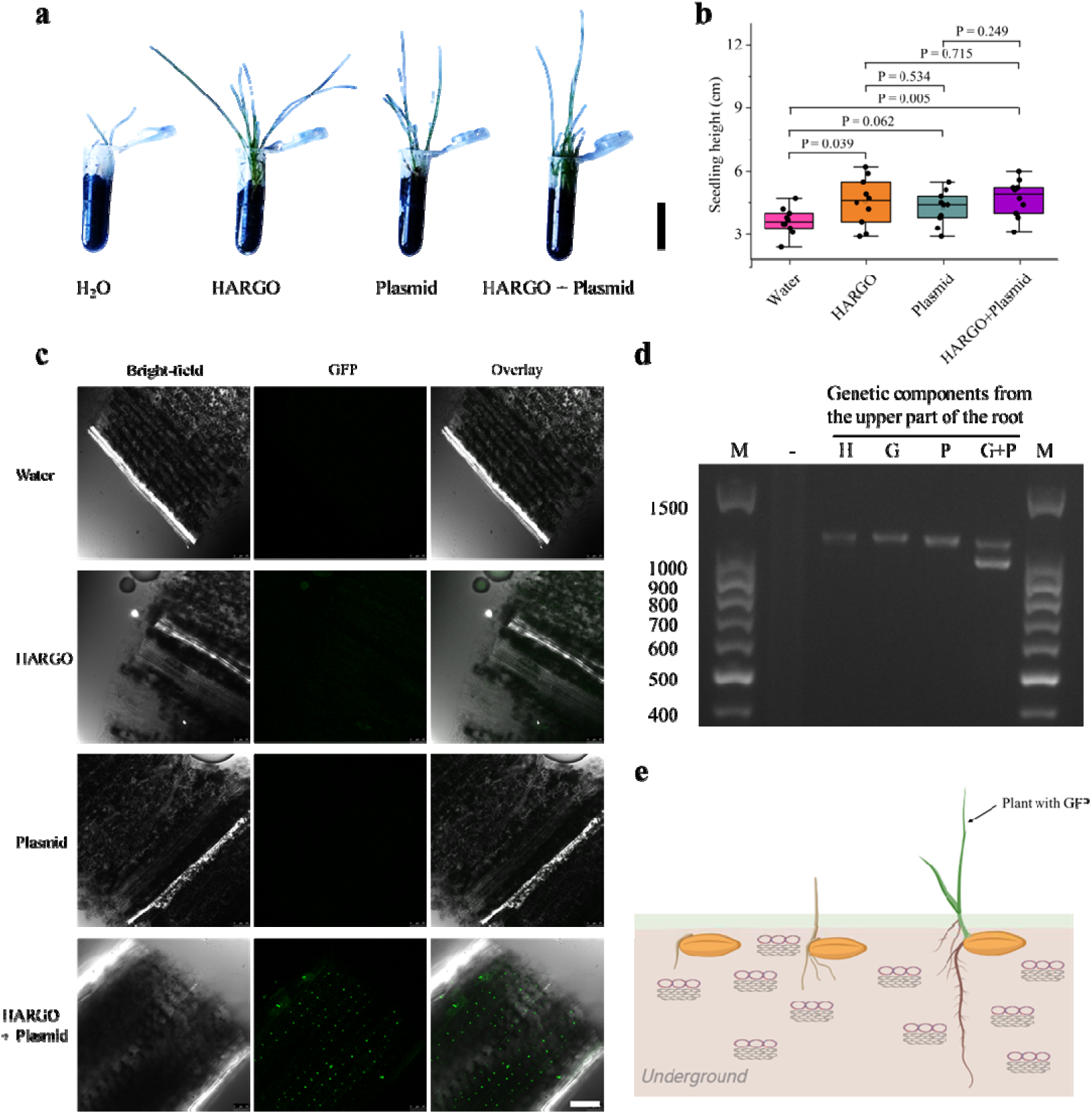
High aspect ratio graphene oxide (HARGO) was used to deliver plasmid DNA pEG100-PcNAC-EGFP to the seeds of *Poa crymophila* Keng and produce positive plants. **a,** A comparison of irrigation using water, HARGO, plasmid DNA pEG100-PcNAC-EGFP, and a mixed aqueous solution of HARGO and pEG100-PcNAC-EGFP, with digital photographs taken 14 days after the emergence of *Poa crymophila* Keng seedlings in dry soil. The scale bar represents 2.5 cm. **b**, Statistical analysis of the seedling lengths of ten randomly selected *Poa crymophila* Keng seedlings. **c**, Laser confocal microscopy images of the leaf tips from four groups of *Poa crymophila* Keng. The scale bar represents 25 µm. **d**, Agarose gel electrophoresis images of PCR products from four groups of *Poa crymophila* Keng seedlings: water (H), HARGO (G), plasmid DNA pEG100-PcNAC-EGFP (P), and a mixed aqueous solution of HARGO and pEG100-PcNAC-EGFP (G+P). **e**, Schematic representation of HARGO delivering plasmid DNA to plant seeds and resulting in positive seedlings.

Subsequently, we performed frozen sectioning on the apical portions of the seedlings, followed by immediate laser confocal detection. The results indicated that no positive signals of green fluorescent protein (GFP) were detected in the Water, HARGO, and Plasmid groups. In contrast, the HARGO + Plasmid group exhibited positive GFP signals, which were predominantly localized in the nuclei and chloroplasts of the apical cells (Fig. 4c). This observation can be attributed to the fact that *PcNAC* is a transcription factor. The localization and function of *PcNAC* within cells dictate that the PcNAC-GFP fusion protein, delivered via plasmid DNA, is predominantly found in the nuclei and chloroplasts. To further verify the delivery of plasmid DNA pEG100-PcNAC-EGFP to *Poa crymophila* Keng cv. Qinghai by HARGO, we conducted PCR analysis on seedlings from the four treatment groups. Consequently, we successfully amplified a specific band corresponding to pEG100-PcNAC-EGFP in the seedlings from the HARGO + Plasmid group, measuring 978 bp, which indicates the presence of plasmid DNA pEG100-PcNAC-EGFP in the seedlings of *Poa crymophila* Keng cv. Qinghai. Finally, to assess the success rate of HARGO in delivering plasmid DNA, we randomly performed cryosectioning and laser confocal detection on the apical parts of 10 *Poa crymophila Keng* cv. Qinghai seedlings. We found that 9 seedlings exhibited positive GFP signals, indicating a 90% success rate for HARGO in plasmid DNA delivery (Fig. S3).

We utilized spring wheat Zhongkemai 1816 (ZKM1816) and rice Zhonghua 11 (ZH11) as experimental materials for further verification. The results indicated that, compared to the control group irrigated with water, the wheat seeds in the experimental group treated with a mixed aqueous solution of HARGO and pEG100-PcNAC-EGFP failed to germinate, whereas the rice exhibited a significant reduction in plant height, demonstrating a pronounced inhibitory effect. These findings suggest that HARGO may effectively deliver pEG100-PcNAC-EGFP to wheat and rice seeds, exerting a significant inhibitory effect on their germination and seedling growth. This suggests that when the pEG100-PcNAC-EGFP plasmid DNA, intended for expression in *Poa crymophila* Keng, enters wheat and rice seeds, it exerts a significant inhibitory effect on seedling emergence (Fig. S4a).

Furthermore, during cryosectioning and laser confocal detection of the apical region of rice seedlings, GFP signals were not detected, suggesting that the promoter associated with pEG100-PcNAC-EGFP is unable to initiate transcription in rice (results not shown). Subsequently, PCR analysis was conducted on the aerial tissues of rice seedlings. The results indicated that no bands were observed in the aerial tissues of rice seedlings from the control groups, whereas a specific PCR band measuring 978 bp was detected in the aerial tissues of rice seedlings from the HARGO + Plasmid group, demonstrating that HARGO effectively delivered plasmid DNA pEG100-PcNAC-EGFP to rice seeds (Fig. S4b).

## Discussion

### HARGO is a novel form of GO

Conventional methods for preparing graphene oxide (GO), including the Brodie, Staudenmaier, Hummers, and modified Hummers methods, typically involve the strong acid oxidation of graphite and consist of three primary steps: intercalation, oxidation, and exfoliation^24^. The thickness of GO sheets produced by these methods is approximately 1 nm, and their morphology typically consists of irregular polygons^24,27^. Electrochemical oxidation is another effective method for preparing GO. Historically, most researchers have concentrated on rapidly preparing GO via electrochemistry; however, few have utilized low-voltage, low-current electrochemical oxidation for extended periods ^28,29,30^.

This study employs an innovative method for preparing graphene oxide (GO) through the electrochemical oxidation of a NaOH aqueous solution at low voltage and current over an extended duration. Although more time-consuming, this approach yields thin-layer GO that is predominantly single-layered and exhibits an elongated strip-like morphology compared to traditional methods. This morphology may result from the uniform oxidation of graphite under low voltage and current conditions, where the current flows from top to bottom along the grinding rod, leading to the characteristic strip shape of GO with a multi-rift structure. The HARGO synthesized in this study demonstrates unique physicochemical properties compared to GO prepared by traditional methods, such as the modified Hummers method and rapid electrochemical oxidation method^24,28^. The strip shape and high aspect ratio of HARGO provide significant advantages for potential applications in biomedicine and materials science. Moreover, the high purity and single-layer structure of HARGO ensure both safety and functionality in nanotechnology applications.

In summary, this study successfully developed a novel type of graphene oxide (GO) through low-voltage, low-current, and longtime electrochemical oxidation. Compared to GO prepared by traditional methods, HARGO exhibits reduced thickness, more uniform morphology, and higher purity, thereby opening new possibilities for its application in various fields.

### The unique physicochemical properties of HARGO contribute to its high penetrability into plant tissues

For a nanomaterial to achieve high penetrability into various plant tissues, it typically requires suitable size (including small particle size and uniform size distribution), favorable surface characteristics (including appropriate surface charge, good hydrophilicity or hydrophobicity, as well as surface modification and functionalization), adequate dispersibility, good biocompatibility, and high stability^31^. In this study, we innovatively synthesized HARGO using a low-voltage, low-current, and long-duration electrochemical oxidation method. HARGO exhibits a distinctive morphological structure. Its aspect ratio is approximately 8:1, with an average thickness of 0.414 nm, presenting a thin, elongated, ribbon-like morphology. Regarding its surface chemical properties, the oxygen-containing group content per unit area is relatively low, while the surface charge per unit mass is high, with concentrations reaching 10 mg/mL or even higher. After being stored at room temperature for 3 years, there were no signs of precipitation or π-π stacking (indicating almost no change in color), fully demonstrating that HARGO possesses favorable surface characteristics, high dispersibility, and remarkable stability. Furthermore, in this study, HARGO not only exhibits no biological toxicity toward plant growth but also promotes plant growth to some extent, demonstrating good biocompatibility.

Based on the unique physicochemical properties of HARGO described above, its high aspect ratio and thin morphology may facilitate its passage through structures such as gaps between plant cells, micropores in cell walls, and channels in cell membranes from a mechanistic perspective. For instance, carbon nanotubes and silicon carbide whiskers, which possess high aspect ratio morphologies, also exhibit notable penetrability in plant tissues^20,18^. Secondly, an appropriate surface charge can interact with the cell membranes of plant cells, promoting the adhesion and entry of HARGO. Furthermore, good hydrophilicity and hydrophobicity (corresponding to the sp^3^ and sp^2^ regions on the surface of HARGO, respectively) enable better adaptation to and penetration of specific tissues based on the distinct characteristics of various plant tissues. High dispersibility ensures that HARGO uniformly contacts plant tissues, while high stability guarantees that its structure and performance remain stable during the process of penetration into plant tissues. In conclusion, HARGO demonstrates excellent penetrability in various plant tissues, particularly in wheat seeds.

### HARGO physically adsorbs plasmid DNA and facilitates its delivery with high penetrability in plant tissues via passive transport

It is widely reported that GO can effectively adsorb plasmid DNA. The specific adsorption mechanism encompasses several physicochemical properties, including local electrostatic attraction, hydrogen bonding between carboxyl, hydroxyl, and epoxy groups with the amino group of plasmid DNA, van der Waals interactions, flexible wrapping of plasmid DNA by GO, and π-π stacking between the bases in GO and plasmid DNA^32,33^. In this study, we employed UV-vis spectroscopy, PCR, and AFM to confirm that HARGO adsorbs plasmid DNA, suggesting that HARGO, similar to conventional GO, can effectively adsorb plasmid DNA through physical adsorption.

Although GO can adsorb plasmid DNA, it cannot directly deliver the DNA to plant cells or various plant tissues. To enable GO to deliver plasmid DNA to plant cells, it must be modified with PEG—primarily to enhance biocompatibility—and PEI—mainly to improve the physical adsorption between GO and plasmid DNA^26^. However, since both PEG and PEI are macromolecules, the physical volume of the GO-PEG-PEI complex is substantial, hindering its ability to deliver plasmid DNA to the deeper layers of plant tissues—only allowing delivery to the surface layers.^26^. In this study, HARGO effectively adsorbs plasmid DNA through physical adsorption without the need for PEG and PEI functionalization (Fig. 3). After adsorbing plasmid DNA, HARGO can also penetrate plant cells and reach deeper layers of plant tissues via passive transport (Fig. S5). This is likely attributed to the small volume of the HARGO-plasmid DNA complex. Additionally, it may benefit from the elongated morphology of the HARGO-plasmid DNA complex, analogous to that of carbon nanotubes and silicon carbide whiskers^20,18^.

## Summary and prospect

This study successfully developed a novel type of graphene oxide, termed HARGO, using a low-voltage, low-current, long-duration electrochemical oxidation method. HARGO demonstrates significant penetration into plant tissues, efficiently adsorbing and delivering plasmid DNA to *Poa crymophila* Keng seeds, leading to the production of transgenic plants with a transformation efficiency of up to 90%. Additionally, HARGO is also effective for wheat and rice.

Compared to traditional plasmid DNA delivery methods—such as Agrobacterium-mediated transfer, gene guns, electroporation, and plant viral delivery methods—HARGO serves as a delivery carrier that offers distinct advantages, including high speed, simplicity, safety, high transformation efficiency, and low cost. Given its unique properties, HARGO is anticipated to play a crucial role in areas such as plant transgenesis, gene editing, and gene knockout, thereby facilitating crop improvement and enhancing agricultural production. Furthermore, considering HARGO’s capability to penetrate tissue cells effectively, it holds significant potential in the medical field and may be employed for tumor-targeted therapy, treatment of genetic diseases, and gene therapy.

Ultimately, the high aspect ratio and penetration capability of HARGO present new opportunities for its development as a nanomedicine carrier, particularly in improving drug delivery efficiency and targeting precision. In conclusion, this study has successfully synthesized GO with a high aspect ratio and demonstrated its potential applications in plant genetic transformation. As a novel class of nanocarriers, HARGO holds significant promise for future applications in biotechnology and medicine, warranting further research and development.

## Declarations

## Acknowledgements

We extend our sincere gratitude to Professor Wang Tao and Professor Fan Xiaoli from the Chengdu Institute of Biology, Chinese Academy of Sciences, for providing the research platform that established a solid foundation for our study. We thank Professor Luo Peigao from Sichuan Agricultural University for his financial support of the initial concept of this research. We also wish to express our special thanks to Professor Ma Xinrong for his in-depth guidance and invaluable support in the field of plant genetic engineering, which proved crucial to our research. Furthermore, we are profoundly grateful to Professor Li Hui for providing the rice seeds (Zhonghua 11), which were pivotal to the experimental aspect of our study. Finally, we thank Professor Yuan Gang from the Department of Thoracic Surgery and the Thoracic Oncology Institute at West China Hospital, Sichuan University, for supplying the essential plasmid extraction reagents crucial for the seamless progression of our experiments.

## Author contributions

C.J. was primarily responsible for the data analysis in this study and took the lead in manuscript preparation. Y.P. made substantial contributions in cryosectioning technology and laser confocal microscopy. Z.Y. successfully prepared and characterized graphene oxide with a high aspect ratio, thereby establishing a foundation for the investigation of material properties. X.L. accomplished the critical experimental steps in plasmid construction, providing a crucial tool for gene function studies. All authors participated in extensive discussions and multiple revisions throughout the manuscript preparation, ensuring the quality and accuracy of the research findings.

## Ethics approval

Not applicable.

## Code availability

Not applicable.

## Availability of data and materials

All data and materials described in this paper are available from the corresponding author upon request.

## Methods

### Plant materials

The variety of *Poa crymophila* Keng is *Poa crymophila* Keng cv. Qinghai (The seed was provided by Professor Ma Xinrong of the Chengdu Institute of Biology, Chinese Academy of Sciences). The wheat variety is spring wheat Zhongkemai 1816 (ZKM1816) and Zhongkemai 138 (ZKM138), and the seeds of them was provided by Professor Wang Tao of the Chengdu Institute of Biology, Chinese Academy of Sciences. The rice variety is Zhonghua 11 (ZH11, the seed was provided by Professor Li Hui of the Chengdu Institute of Biology, Chinese Academy of Sciences).

### Preparation of HARGO

Graphite rods with a 5 mm diameter (spectral purity, model YL-BDC-100) were used as the anode, while a Pt electrode of 99.99% purity (10*10*0.1 mm) served as the cathode. In a 400 mL solution of 0.5 mol/L NaOH, the electrode spacing was set at 6.5 cm, with both electrodes submerged to a depth of 6.5 cm in the electrolyte. Electrochemical oxidation was carried out at a voltage of 3.5 V and a current of 0.03 A for 96 h. The resulting brownish-yellow solution after electrolysis was dialyzed against deionized water using MD44-3500 dialysis tubing until the pH stabilized at 6.8. The solution was then concentrated at a constant temperature of 40_ in an oven after dialysis. The HARGO concentration was determined by drying 100 mL of the concentrated solution to a constant weight.

### Apparatus for characterizations

The Zeta potential was measured by Nano9200 nanoparticle analyzer (Haixinrui). Atomic force microscopy (AFM) images of the samples were captured in tapping mode using a Nano Scope IIIA AFM, with samples prepared by spin coating a diluted aqueous solution onto a mica substrate at 1000 r.p.m. Fourier transform infrared spectroscopy (FT-IR) spectra were recorded on a PE Spectrum 100 spectrometer, utilizing either a film or KBr discs. Ultraviolet-visible spectroscopy (UV-Vis) spectra were obtained using a PerkinElmer Lambda 35 UV/Vis Spectrometer for analysis in the ultraviolet-visible range.

### Titration test

For the HARGO group, 10 mg of HARGO was dissolved in 25 mL of 0.1 mol/L HCl aqueous solution (adjusted to pH = 1.08), and then titrated with 0.1 mol/L NaOH aqueous solution. For the control group, 25 mL of 0.1 mol/L HCl aqueous solution (adjusted to pH = 1.08) was taken and titrated with 0.1 mol/L NaOH aqueous solution. During the titration process, the pH value change of the solution was detected in real time by a pH meter (PHS-3E, Shanghai, China). The titration curve was plotted by Origin v2021 software.

### Construction of plasmid DNA pEG100-PcNAC-EGFP

Took 20 µL of plasmid DNA backbone pEG100-EGFP with a concentration of 100 ng/µL. Then, perform enzymatic digestion with BamH _ and BbvC _ according to the instructions. Subsequently, performed agarose gel electrophoresis and gel recovery to obtain the digested pEG100-EGFP. Used the Plant Total RNA Extraction Kit (BL1180A) to extract the total RNA of *Poa crymophila* Keng, and then detect the concentration and quality of the extracted total RNA by agarose gel electrophoresis. Used the reverse transcription kit (Cat. No. 218161) to synthesize cDNA using RNA as a template. Performed PCR amplification with specific primers and sequencing to obtain a *PcNAC*-specific fragment with one intron deletion. The upstream sequence of the specific primer is 5’-GAATTCATGGGGATGGCCGTGCGCAGG-3’, and the downstream sequence is 5 ‘ -CCTCAGCGGAGTCGCTCAAGAAGGGAGCCGGCATGCC-3 ’ . Then, performed PCR amplification on this specific fragment using primers with restriction endonuclease-specific sequences of BamH _ and BbvC _ added at both ends, and performed agarose gel electrophoresis and gel recovery to obtain the target DNA fragment. Add the obtained target DNA fragment aqueous solution (volume 20 µL, concentration 100 ng/µL) to restriction endonucleases BamH _ and BbvC _ for enzymatic digestion. Then, performed agarose gel electrophoresis and gel recovery again to obtain the digested target DNA fragment. Use the T4 DNA ligase kit (Cat. No. M02002) to ligate the digested plasmid DNA pEG100-EGFP and the target DNA fragment to obtain plasmid DNA, specifically pEG100-PcNAC-EGFP. The length of the plasmid DNA is 11, 798 bp.

### Adsorption of plasmid DNA onto HARGO

The constructed pEG100-PcNAC-EGFP was transformed into DH5 α *Escherichia coli* and cultured on LB solid medium containing kanamycin. After inverted culture at 37°C for 2 days, a single colony was picked from the plate and inoculated into liquid medium containing kanamycin, and cultured overnight with shaking at 200 rpm at 37°C to make the bacteria grow to the late logarithmic growth phase. The cultured bacterial solution was transferred to a centrifuge tube. After centrifugation at 4000 rpm for 10 minutes, the supernatant was discarded and the bacterial cell pellet was collected. Subsequently, plasmid DNA was extracted according to the instructions of QIAGEN Plasmid Maxi Kit (Cat. No. 12163), and the extracted plasmid DNA was quality tested and quantified by NanoDrop 2000/2000c (NanoDrop Technologies; Thermo Fisher Scientific, Inc., Pittsburgh, PA, USA).

A mixture was prepared by combining a 0.5 mg/mL aqueous solution of HARGO with a 110 ng/µL aqueous solution of plasmid DNA (pEG100-PcNAC-EGFP, 11,798 bp in length) in equal volumes. The components were gently agitated at room temperature to ensure thorough mixing, yielding an aqueous solution of HARGO with adsorbed plasmid DNA.

### Delivery of plasmid DNA to plant seeds via co-cultivation with HARGO

In 2.0-mL polyethylene (PE) tubes, a small amount of dry substrate soil was placed, *Poa crymophila* Keng seeds were sown, and covered with a thin layer of dry substrate. The soil was then irrigated with the aqueous solution of HARGO adsorbed with plasmid DNA, ensuring that the soil was moist with the solution level about 1 mm above the soil surface. The tubes were kept in darkness at room temperature for two days to facilitate the delivery of plasmid DNA to the seeds by HARGO. Following the dark treatment, the plants were transferred to a phytotron with a photoperiod of 12 hours light/12 hours darkness, a light intensity of 300 photosynthetic photon flux density (PPFD), a temperature of 22-25_ during the light hours, and 18-20_ during the dark hours.

### Laser confocal microscopy detection

Leaf sections from 14-day-old seedlings were obtained, and 100 µm thick transverse sections were cut using a cryostat (Leica CM1950). These sections were promptly examined under a laser confocal microscope (Leica TCS SP8), using 488-nanometer excitation light and a detection wavelength range of 500-540 nanometers.

### PCR analysis

Stems and leaves of *Poa crymophila* Keng from 14-day-old seedlings were sampled and processed for PCR amplification following the instructions provided in the Plant Leaf Direct PCR Kit (Cat. No: TP-0211T). The PCR reaction mixture consisted of 20 µL, comprised of 10 µL of 2×Leaf PCR Easy^TM^ Mix, 1 µL each of upstream and downstream primer (10 mM), 1 µL of sample lysate, and 7 µL of ddH_2_O. The upstream primer sequence was 5‘-ATGGGGATGGCCGTGCGCAGG-3’, and the downstream primer sequence was 5‘-CTCAAGAAGGGAGCCGGCATGCC-3’. The PCR program included initial denaturation at 95_ for 5 min, followed by 32 cycles of denaturation at 95_ for 40 s, annealing at 67_ for 40 s, extension at 72_ for 1 min, and a final extension at 72_ for 5 min. After PCR, 2 µL of 10×DNA loading buffer was added for agarose gel electrophoresis detection.

### Data compilation and statistical analysis

Data were organized using Microsoft Excel 2023. Statistical analyses were conducted using IBM SPSS v24.0 (SPSS, Chicago, USA).

## Supplemental Figure

**Extended Data Fig. 1.**
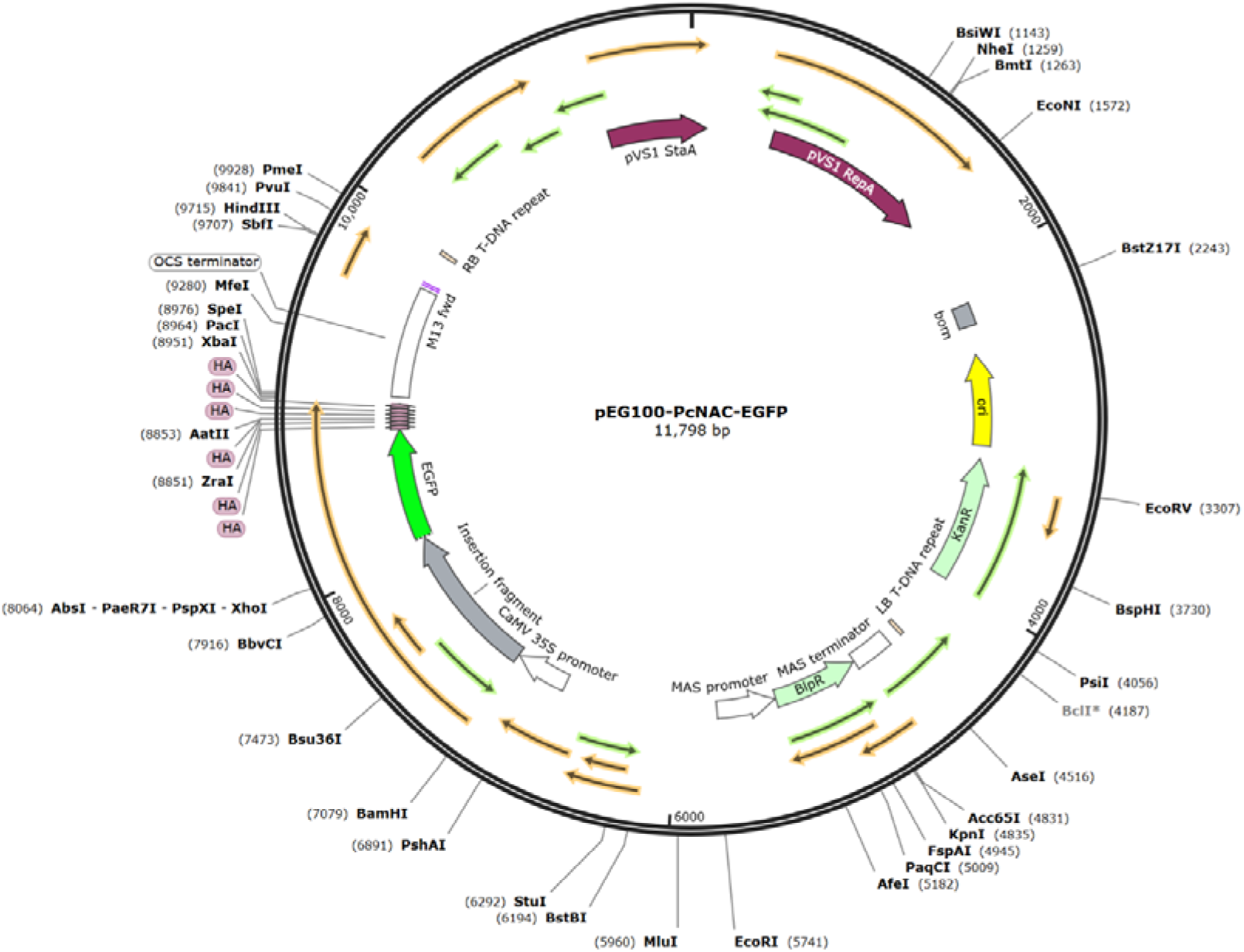
Plasmid map of pEG100-PcNAC-EGFP.

**Extended Data Fig. 2.**
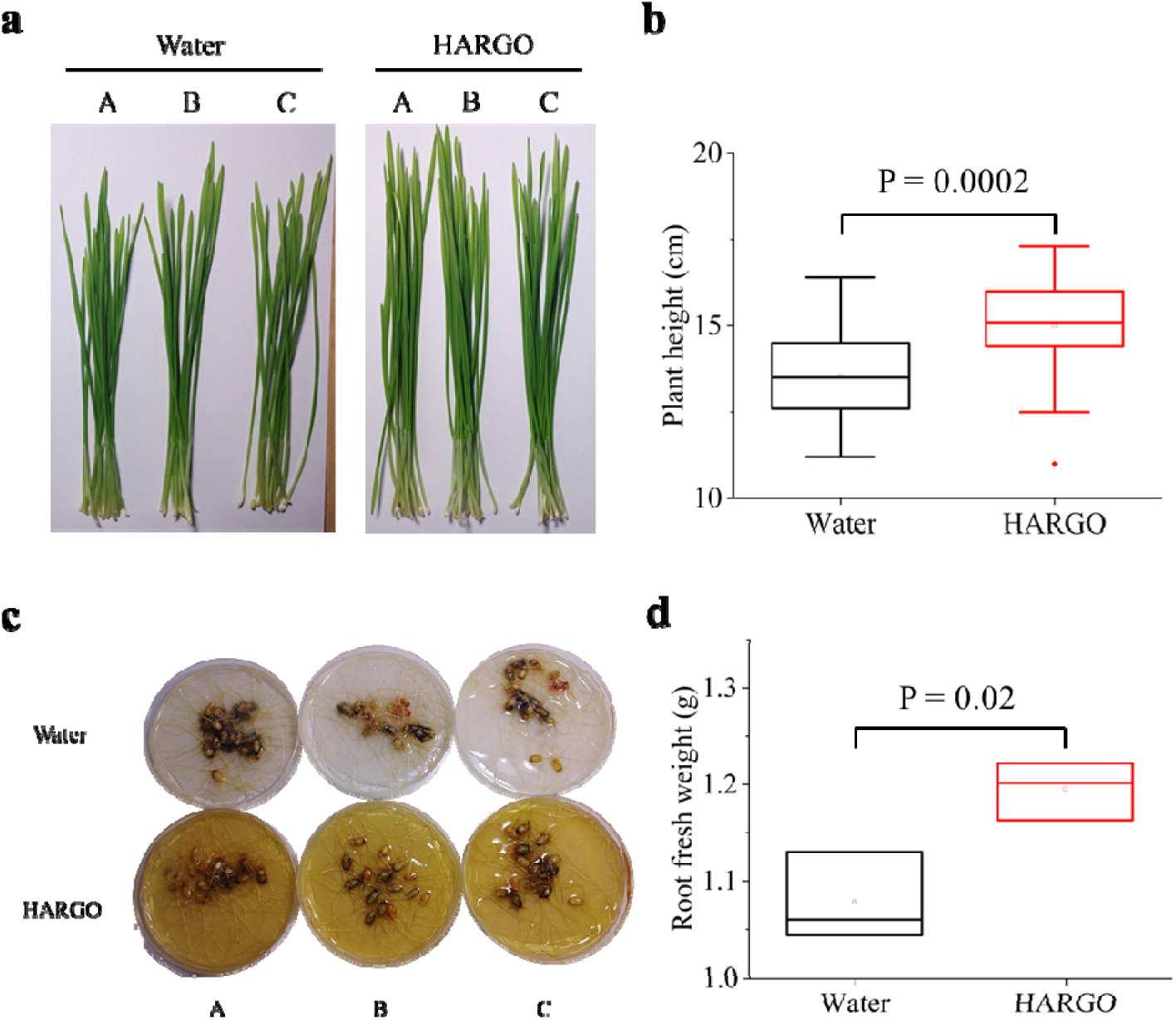
Effect of high aspect ratio graphene oxide (HARGO) on wheat seedlings. a-b, Randomly select 15 seeds of spring wheat variety Zhongkemai 138 (ZKM138) and place them in a culture dish for germination. Digital photo of wheat seedlings 14 days after emergence (a) and their height statistics (b). Among them, the number of seedlings in both groups is 45. **c-d,** Digital photos of the culture dish after 14 days of seedling emergence (c) and statistics of root fresh weight (d).

**Extended Data Fig. 3.**
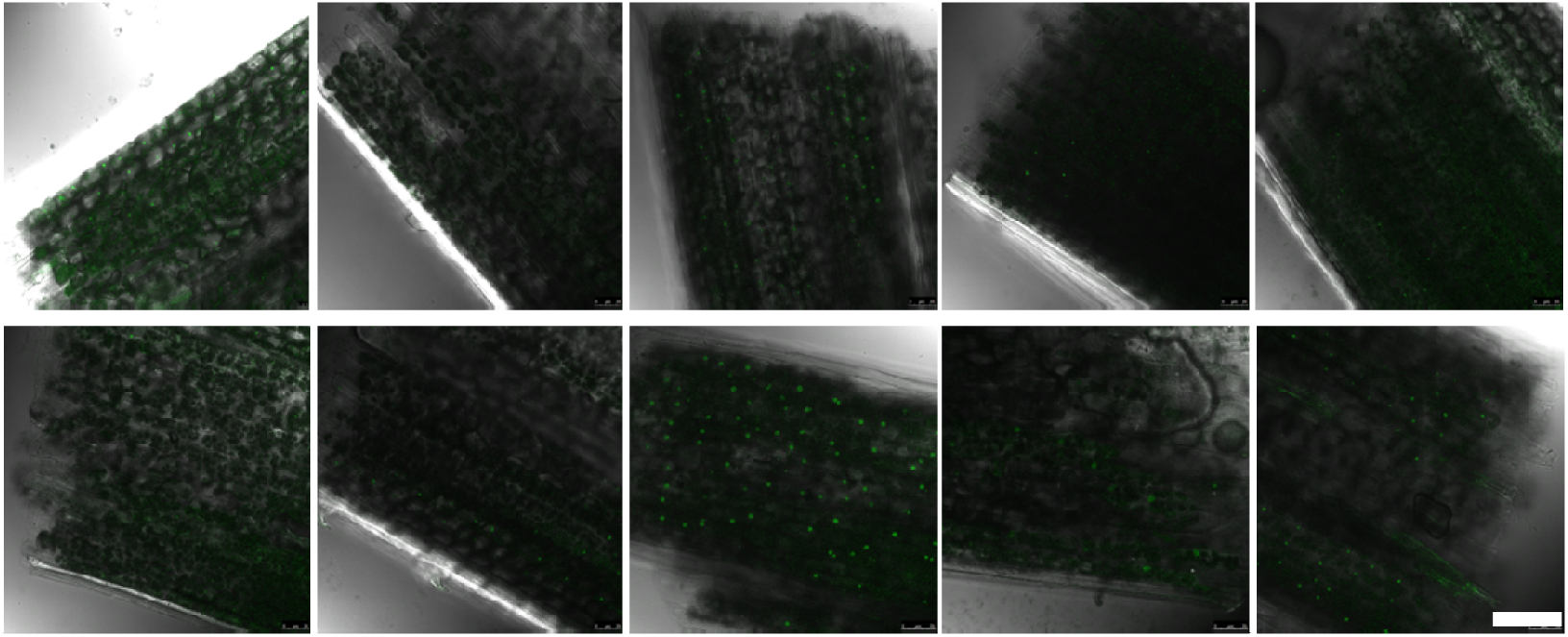
Laser confocal microscopy images of ten 21-day-old *Poa crymophila* Keng seedlings from the HARGO plus plasmid DNA group. The excitation wavelength is 488 nm. The scale bar length is 50 µm.

**Extended Data Fig. 4.**
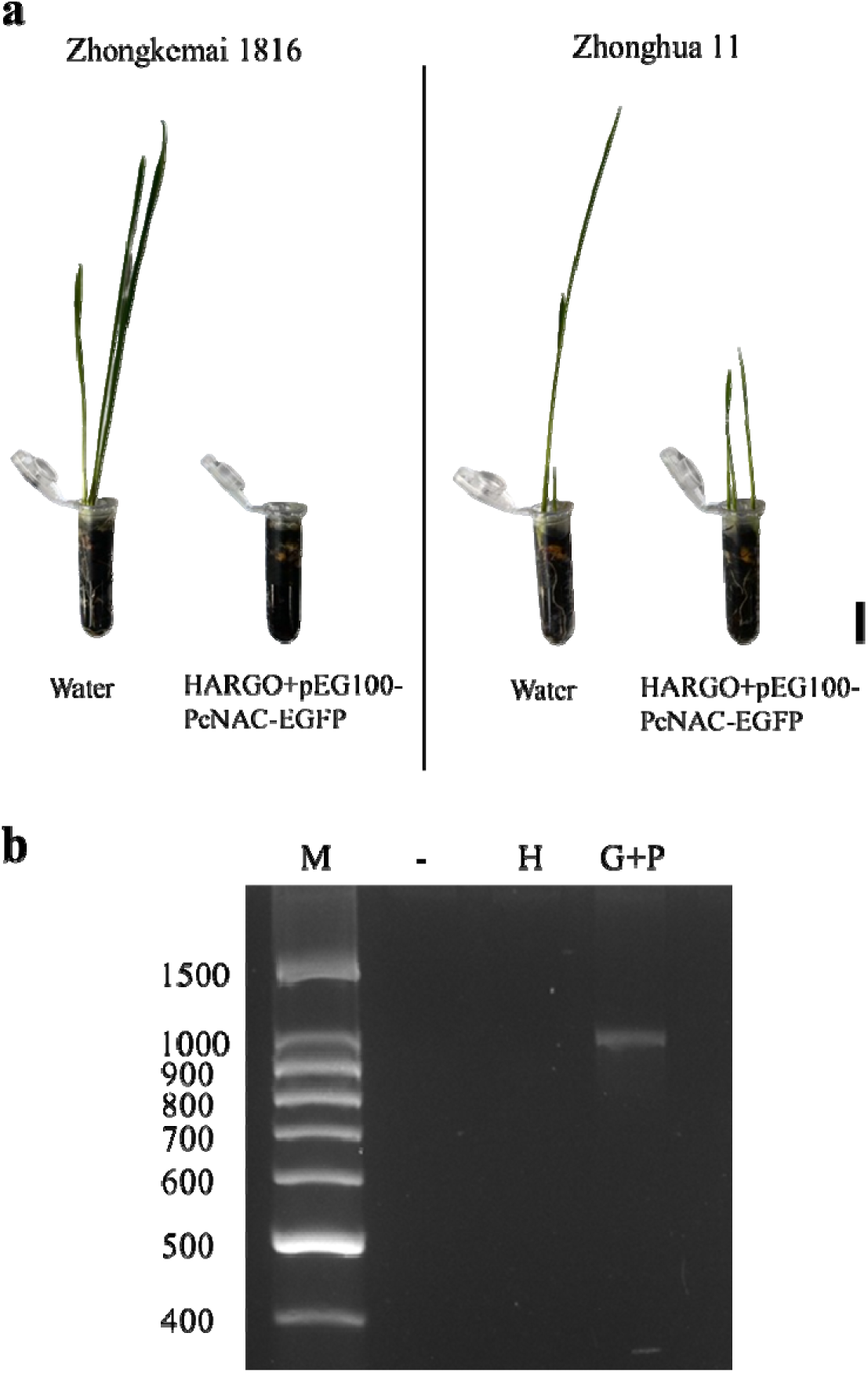
The effect of high aspect ratio graphene oxide (HARGO) delivery of plasmid DNA pEG100-PcNAC-EGFP to wheat and rice seeds on their germination. **a**, Digital photos of spring wheat variety Zhongkemai 1816 (ZKM1816) and rice variety Zhonghua 11 (ZH11) after 10 days of seedling emergence. The scale bar represents 1.5 cm. **b**, Digital photos of agarose gel electrophoresis for PCR detection of rice seedlings.

**Extended Data Fig. 5.**
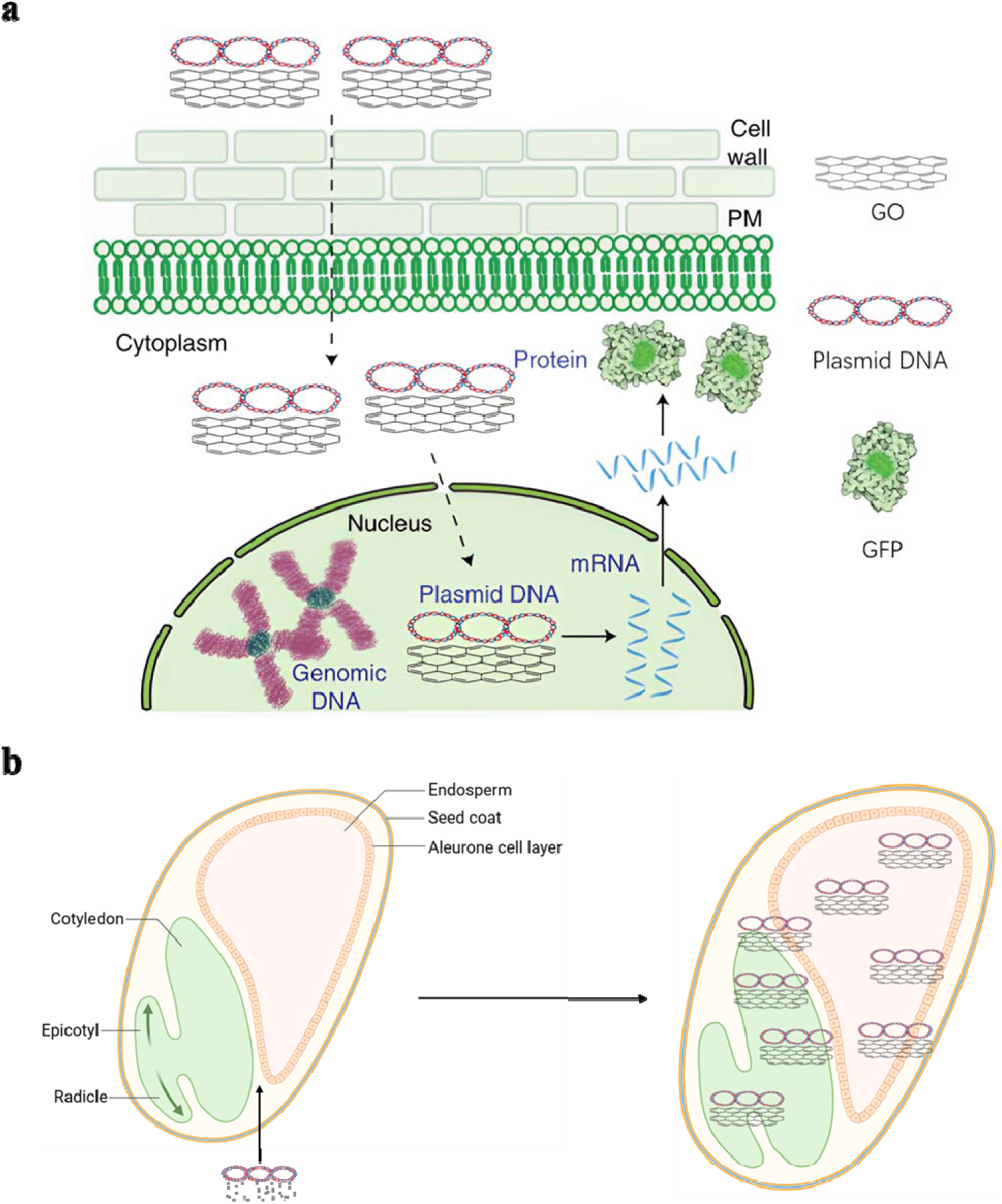
Schematic diagram of HARGO and adsorbed plasmid DNA entering plant cells and various tissues of seeds by passive transport. **a**, After adsorbing plasmid DNA, HARGO passes through the cell wall and cell membrane and enters the cytoplasm, and further enters the nucleus through the nuclear pore. After entering the nucleus, the plasmid DNA is transcribed and translated into mRNA, and the mRNA further enters the cytoplasm through the nuclear pore and is translated into green fluorescent protein. **b**, After adsorbing plasmid DNA, HARGO enters various tissues of the seed with high penetrability.

